# Study of droplet generation in an E-shaped microchannel using two-phase level-set method

**DOI:** 10.1101/2022.11.28.518020

**Authors:** Liana Chatterjee, Sulagna Chatterjee

## Abstract

Droplet-based microfluidics is the study of microfluidic device, where droplets are generated mainly using two methods i.e., active, and passive methods. The manipulation of these droplets in microfluidic channels is an immensely useful technology in various scientific fields such as biological, biomedical studies, and drug delivery. Recently droplets with a core-shell structure have achieved considerable interest due to their unique properties and varied applications.

Droplet splitting as an important feature of droplet-based microfluidic systems has been widely used. In the present paper, two-dimensional numerical simulations have been done to examine the droplet splitting process in an E Junction geometry microchannel. The two-phase level set method (LSM) has been used to analyze the mechanism of droplet formation and droplet splitting in an immiscible liquid/liquid two-phase flow. Governing equations on the flow field have been studied and solved using COMSOL Multiphysics software. The model was developed to simulate the mechanism of droplet splitting at the E-junction microchannel. This study provides a passive technique to split up microdroplets at the E-junction.

## 1. Introduction

Droplet-based microfluidics manipulates discrete volumes of fluids in immiscible phases with low Reynolds number and laminar flow regimes. Studies in droplet-based microfluidics systems have been growing significantly in the previous years. Microdroplets offer the feasibility of handling miniature volumes (μL to fL) of fluids and provide better mixing, encapsulation, sorting, sensing and are also suitable for high throughput experiments. Two immiscible phases used for the droplet-based systems are referred to as the continuous phase i.e., the medium in which droplet is generated, and the dispersed phase which is the droplet phase. Droplet generation originates from fluid instabilities.

Methods for droplet generation are further divided into passive and active, where passive method generates droplets without external stimulus, but the active method makes use of additional energy input for droplet generation. Microfluidic channel provides the boundary of microflow and thus its geometry impacts the droplet generation as well.

Pingan et al. [^1^] presented a unified review of both passive and active droplet generation in a systematic manner from the point of view of physical mechanisms in their study of the two methods of droplet generation where they explained that the Methods for droplet generation can be either passive or active, where the former generates droplets without external actuation, and the latter makes use of additional energy input in promoting interfacial instabilities for droplet generation. In passive method, droplets can be produced in squeezing, dripping, jetting, tip-streaming, and tip-multi-breaking modes, depending on the competition of capillary, viscous, and inertial forces. In active control, droplet generation can be manipulated by either applying additional forces from electrical, magnetic, and centrifugal controls, or modifying intrinsic forces via tuning fluid velocity and material properties including viscosity, interfacial tension, channel wettability, and fluid density. Abrishamkar et al. [^2^] In their work numerically investigated the process of droplet generation using water as the dispersed phase and FC-40 oil as the continuous phase in a microfluidic device. they used a two-phase level set method in COMSOL Multiphysics^®^ for their study. The simulations were carried out in a two-dimensional domain through a flow-focusing droplet generator. Olson et al. [^3^] In their paper, they continued to develop and study the conservative level set method for incompressible two-phase flow with surface tension. Lei et al. [^4^] presented a numerical study of the formation of droplets in a novel two-dimensional T-junction device by using a commercial CFD package: COMSOL Multiphysics. They carried out numerical simulations for different flow conditions and different flow rates which in turn led to four regimes.

In Ich-Long et al. [^5^] paper they presented droplet formations in microfluidic double T-junctions based on a two-dimensional numerical model applying the volume of fluid method. Jayaprakash et al. [^6^] did their study on a 3d T Junction wherein water was the dispersed phase and mineral oil was the continuous phase their paper provides new insights on the pressure difference between the dispersed phase and the continuous phase during the droplet breakup process.

In our study, we have worked with an E-shaped microchannel which is based on cross-flow. The cross-flow geometry refers to the one where dispersed and continuous phase fluids meet at an angle θ (0° < θ ≤ 180°). Furthermore, in such type of microchannels, the two fluids are injected through three channels wherein the continuous phase flows through the upper and lower channel and the dispersed phase through the middle channel, very similar to an “E” the head-on device reduces into an E-junction which is a novel concept reported in this study and whose simulations have further been documented.

## 2. Theoretical Modeling

In this section we describe the construction and development of an E-Junction device in COMSOL Multiphysics. We present the construction of the device (**section 2.1**), the equations used to solve this modelling and flow simulation (**section 2.2**), the conditions set for the simulation (**section 2.3**), the liquids used to flow through the channel (**section 2.4**), and the mesh that was used (**section 2.4**).

### 2.1. Geometry

An E-junction microchannel was created to generate micro-sized droplets. Figure 1 demonstrates the studied channel geometry in this paper which contains three horizontal channels. The continuous phase flows along the upper and lower channel whereas the dispersed phase flows through the middle channel. The measurements of different regions of the E-Junction are given in Table 1 and the figure of the geometry is attached under Figure 1.

**Figure 1:**
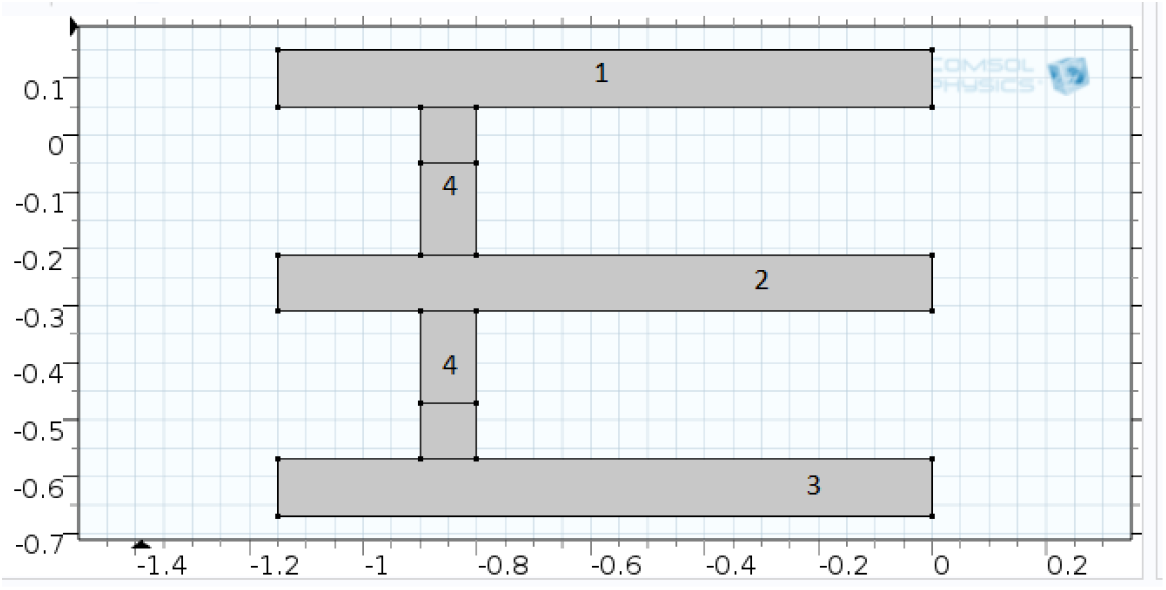
The geometry used in E-Junction simulation

**Table 1:** Measurements of E Junction Geometry

| GEOMETRY | WIDTH | HEIGHT |
| --- | --- | --- |
| RECTANGLE 1,2,3 | 1.15mm | 0.1mm |
| RECTANGLE 4 | 0.1mm | 0.82mm |

### 2.2. Governing Equations

The level set method is an approach that is used to represent moving interfaces or boundaries using a fixed mesh. It is useful for this problem because the level set method is used for simulations where the computational domain can be divided into two domains separated by an interface such as this. In COMSOL Multiphysics, *φ* is a smooth step function that equals zero in one domain and one in the other. Across the interface, there is a smooth transition from zero to one.

In this method, the model comprises the following governing equations.[^3^]

#### Continuity equation

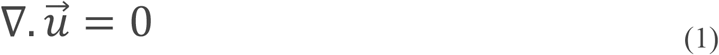

Liquid-liquid two-phase microflows are well described with the continuum hypothesis. The incompressible continuity equation reads, for both dispersed and continuous phase fluids. *∇* is the del operator, *u* is the velocity vector of the fluid.

#### Incompressible Navier-Stokes equation

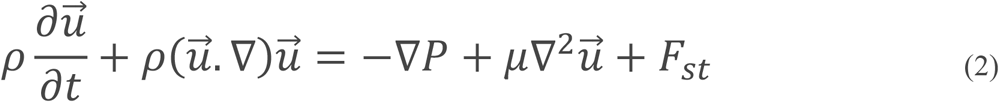

The momentum equation is the well-known Navier-Stokes equation of incompressible Newtonian fluids. where *t* is the time, *ρ* the density of the fluid, *p* the pressure, *η* the dynamic shear viscosity, and *f* the body force vector per unit volume. Also, *F_st_* is the surface tension force, which acts at the interface between two phases.

#### Level-Set equation

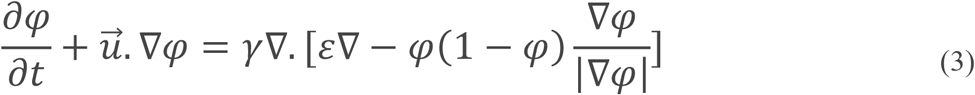

Where *u* is the velocity vector and *t* refers to time. *ρ* and *μ* denote the density and dynamic viscosity, respectively. The pressure is given by *p* while *φ* denotes the level set function. The left-hand terms of the level set equation give the correct motion of the interface, while the right-hand terms are necessary for the stability of the numerical solution. *ε* denotes an artificial thickness of the interface and is typical of the same order as the mesh element size. If *ε* is too small, the thickness of the interface might not remain constant, and oscillations may appear because of numerical instabilities. The parameter *γ* determines the reinitialization or stabilization amount of the *φ*. It needs to be tuned for each specific problem. If *γ* is too large, the interface moves incorrectly. A suitable value for *γ* is the maximum magnitude of the velocity field u.

In this work, the droplet effective diameter (d) is the main output parameter. To obtain the droplet effective diameter an area integration of the volume fraction of the dispersed phase (where φ > 0.5) is performed, and it can be calculated using the following expression:

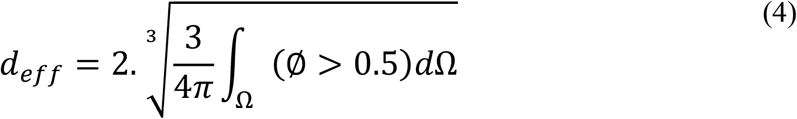

Here, Ω represents the leftmost part of the horizontal channel, where x< −0.7 mm.

### 2.3. Boundary Conditions

At the inlets, the laminar inflow boundary conditions with specified volumetric flow rates as defined under Table 2 are used, the figures are given under Figure 2. The expression is given as step1 under V1, V2, and V3 defines volumetric flow at the inlet boundary using the curve parameter “s”, which is a predefined curve parameter on boundaries in 2D models and runs from 0 to 1 along the direction of the boundary. The smoothed step function ramps up the volumetric flow at the start of the simulation. The step function’s input “t[1/s]” is the predefined variable “t” for the time having unit seconds and the “unit multiplication” using “[1/s]” is done to make the step function’s input dimensionless. At the outlet, the pressure condition is set. The Wetted wall Multiphysics boundary condition applies to all solid boundaries with the contact angle specified as 3*pi/4 radians and a slip length equal to 5e-6 m. The contact angle is the angle between the fluid interface and the solid wall at points where the fluid interface attaches to the wall. The slip length is the distance to the position outside the wall where the extrapolated tangential velocity component is zero.

**Figure 2:**
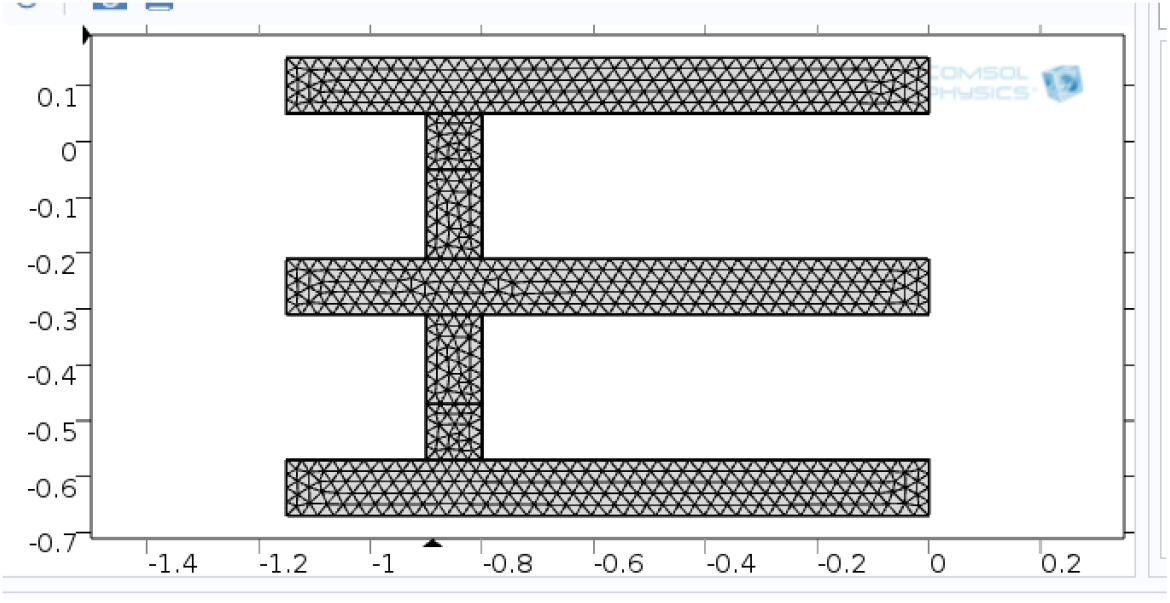
Physics Enabled Mesh being shown.

**Table 2:**
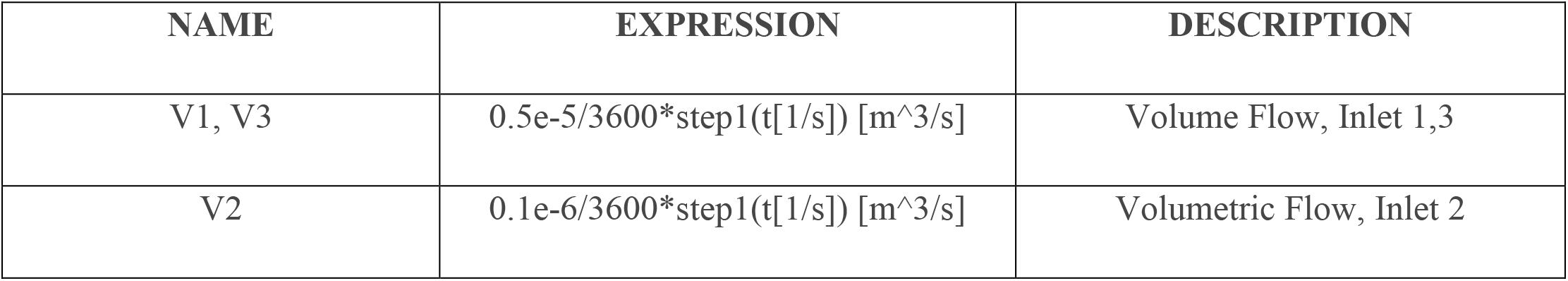
Values of Volumetric Flow rates taken for Inlet 1,2 and 3

| NAME | EXPRESSION | DESCRIPTION |
| --- | --- | --- |
| V1, V3 | $0.5e-5/3600 \cdot \text{step1}(t[1/s])$ [m<sup>3</sup>/s] | Volume Flow, Inlet 1,3 |
| V2 | $0.1e-6/3600 \cdot \text{step1}(t[1/s])$ [m<sup>3</sup>/s] | Volumetric Flow, Inlet 2 |

### 2.4. Fluid Properties

Water and engine Oil have been chosen as working fluids in this study. Water is the continuous phase and engine Oil is the dispersed phase. Physical properties (density and dynamic viscosity) of Water and engine Oil were taken from the built-in materials property package of COMSOL.

### 2.5. Mesh Study

A structured two-dimensional mapped mesh has been generated to discretize the planar geometry of the microchannel in this work. In Figure 2, the grid profile of the E-junction microchannel is illustrated.

## 3. Results & Discussions

### 3.1. Droplet Splitting

Figure 3 is being shown below and attached with it are the time ranges for which droplet splitting took place for this type of geometry. The images gradually begin from 0s and move till 0.26s to show the droplet splitting of oil in water in both the channels of the continuous phase.

Besides the image of the droplet splitting the different processes of the splitting phenomenon has also been mentioned.

**3.**
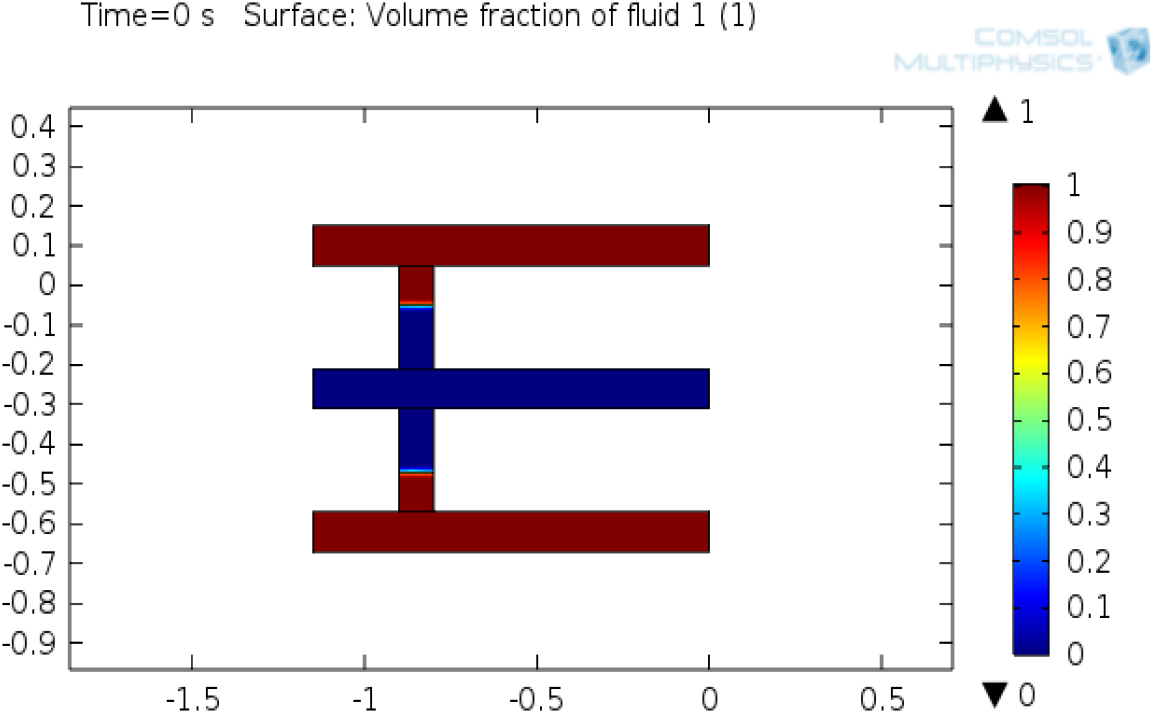
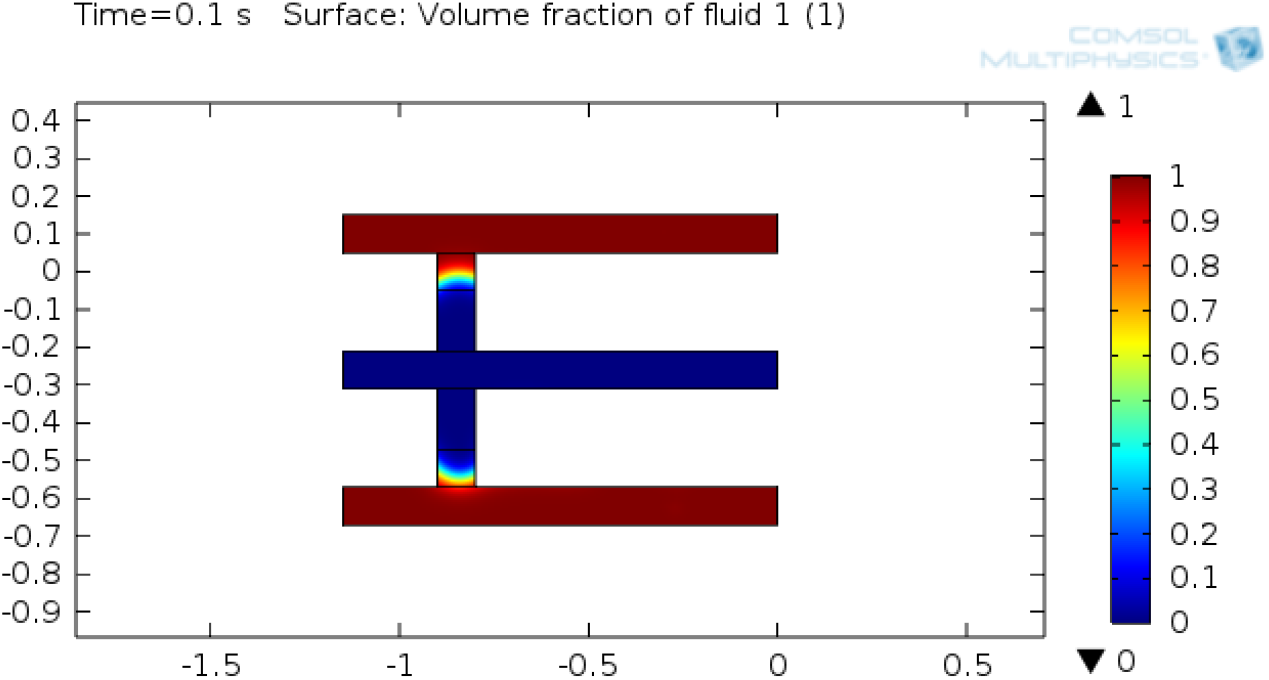
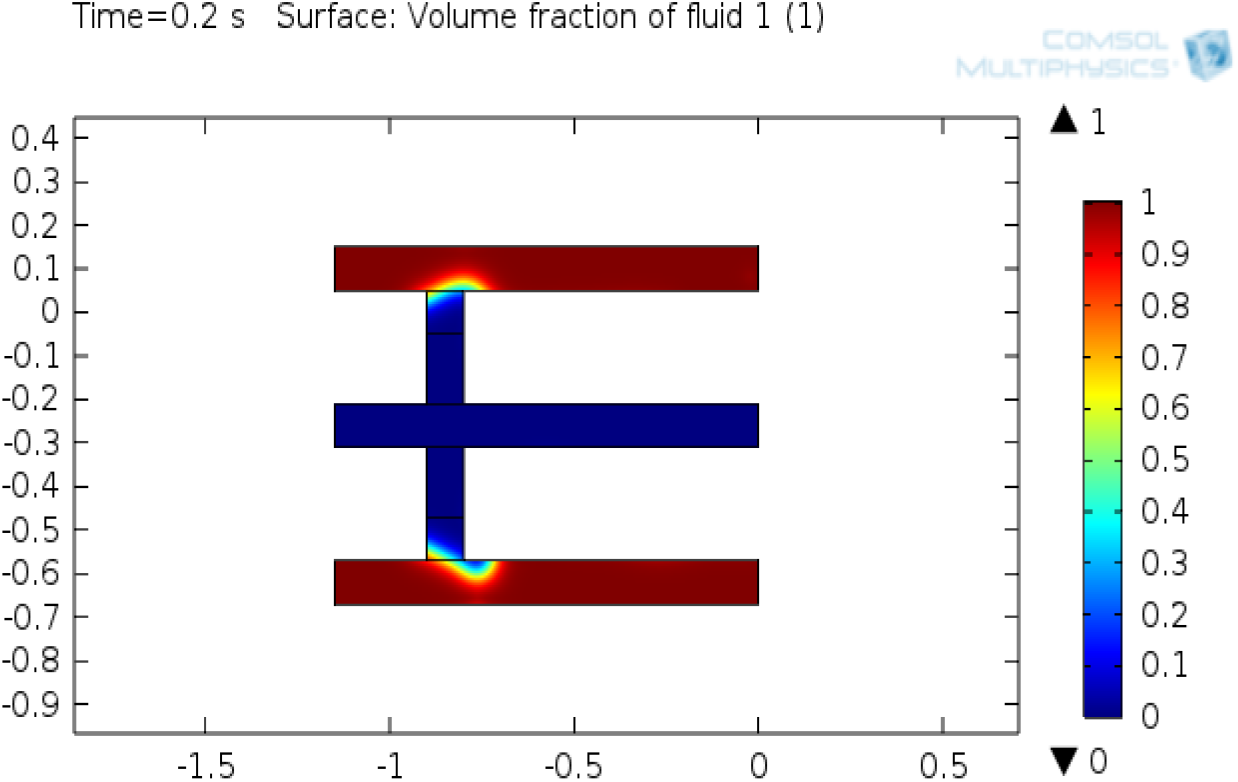
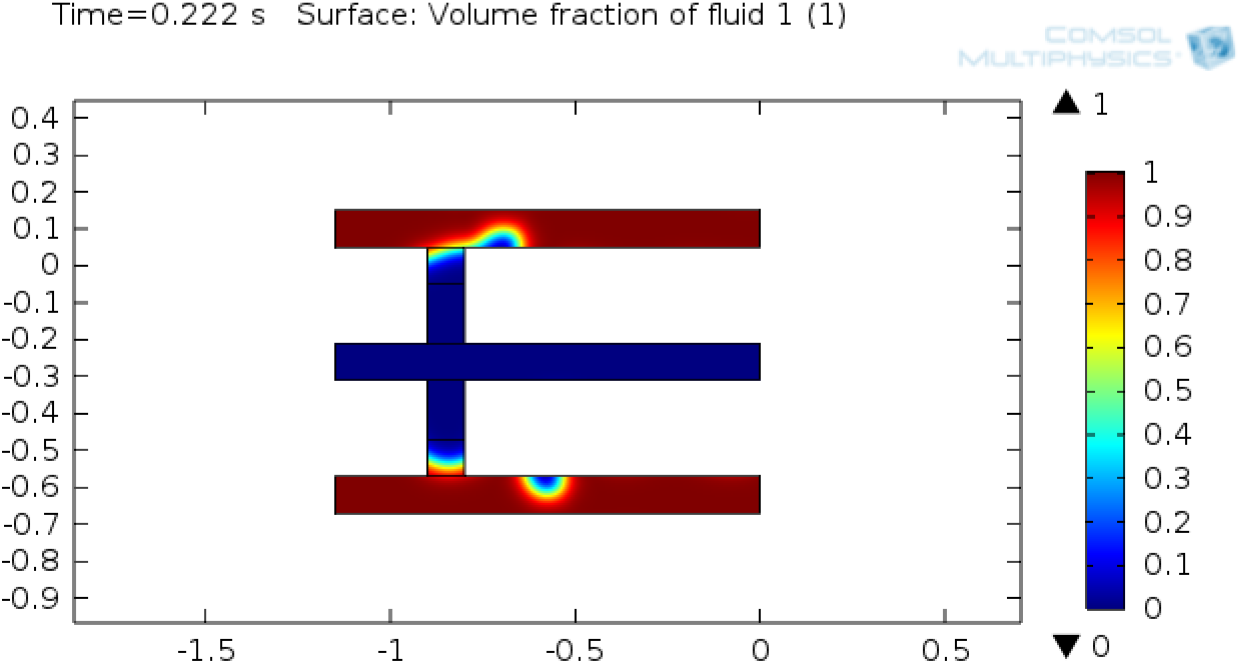
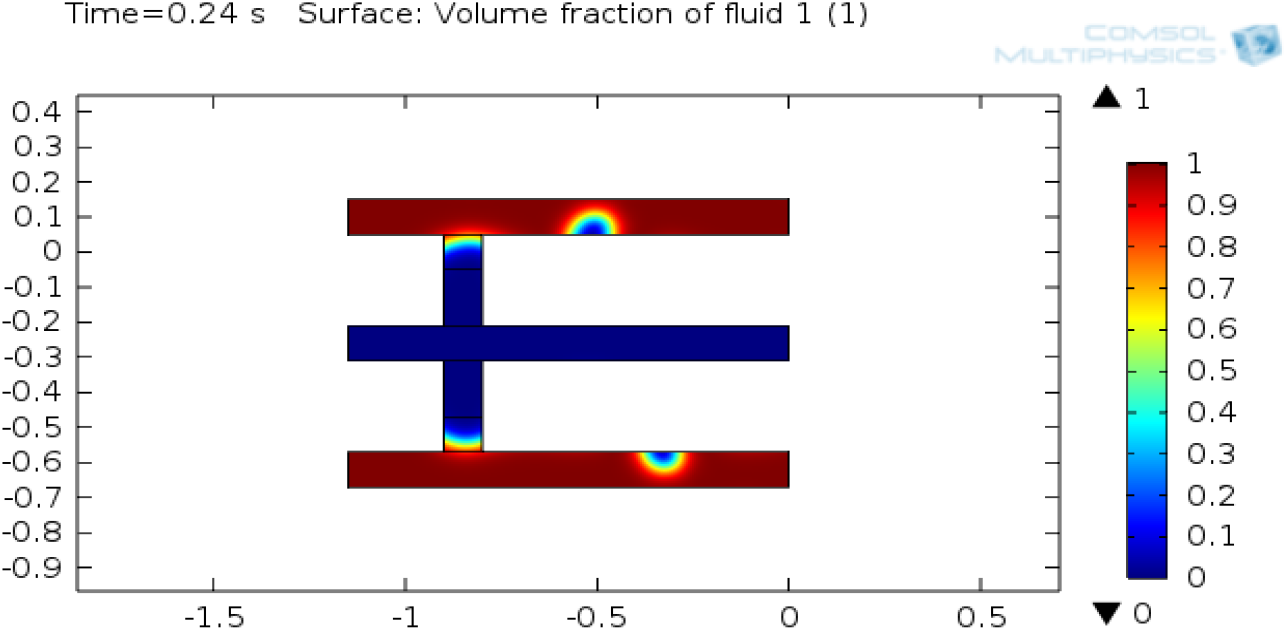
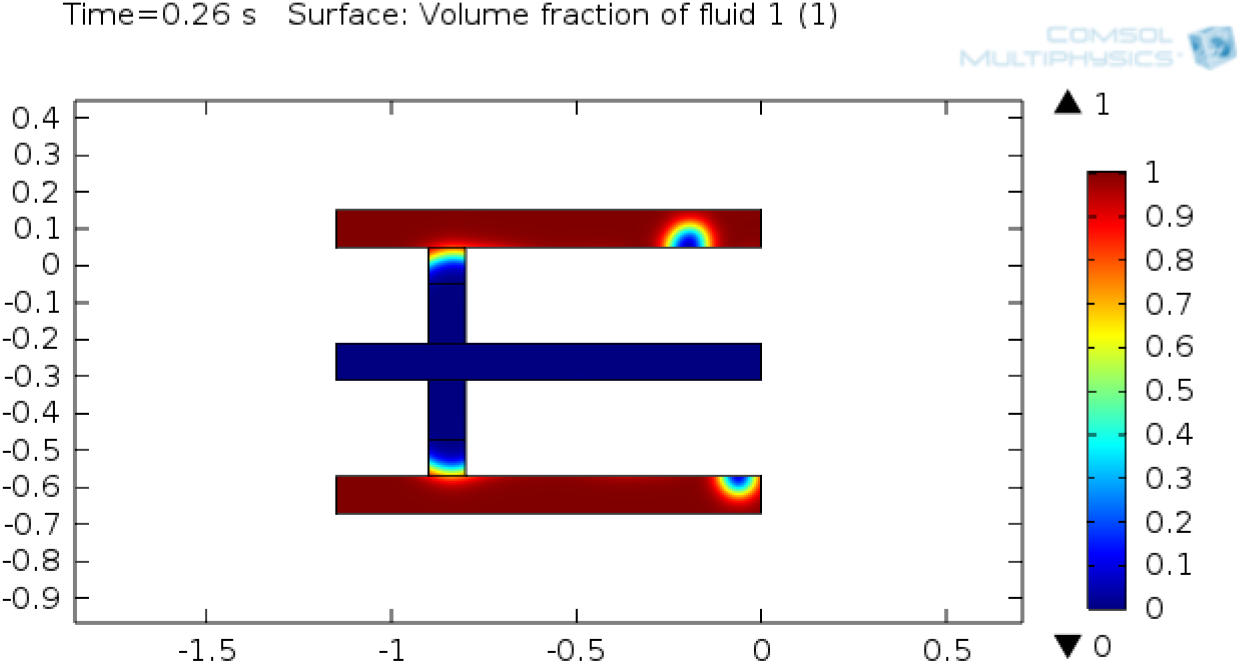
*(i)*Start of the process at t=0s. *(ii)*Filling taking place at t=0.1s *(iii)*Necking process taking place at t=0.2s. *(iv)*Pinch off being shown at t=0.222s. *(v)*Droplets after splitting being shown at t=0.24s. *(vi)*Droplet splitting being shown at t=0.26s.

### 3.2. Study of effective droplet diameter

Upon generating a graph of deff vs time which has been added below it is evident that there is a gradual rise beginning from the pinch-off process that is when a droplet is visible in the channels. It should be noted that during the filling and necking process there is no change in the graph because the effective diameter of the droplet will only be counted once droplet splitting begins. The graph attached under Figure 4 validates this.

**Figure 4:**
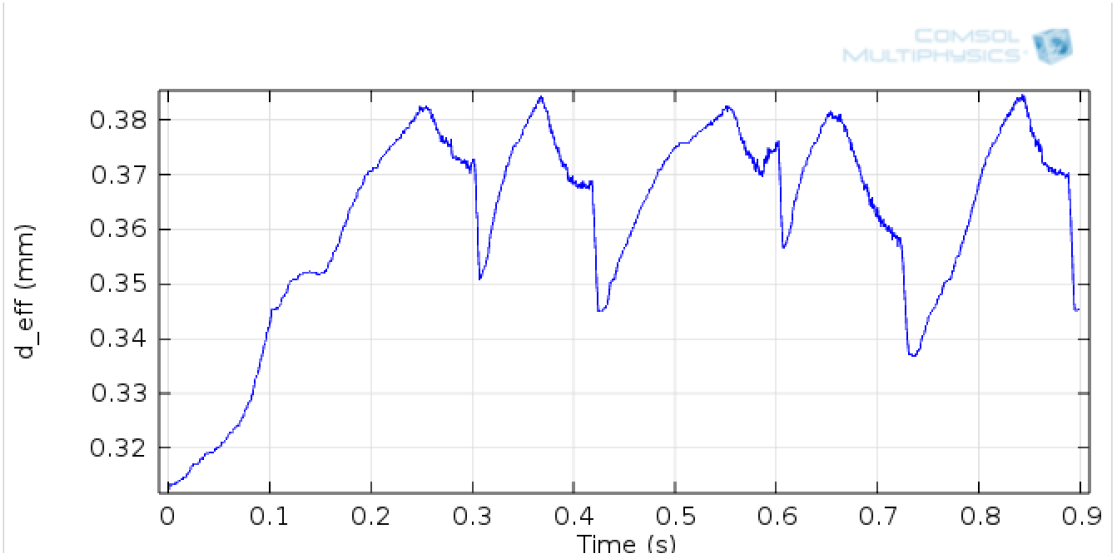
Effective droplet diameter(mm) vs time(sec).

### 3.2. Study of velocity profile

The velocity profile for the respected processes mentioned in section 3.1 have been attached further in Figure 5 which indicates that there is no change in the velocity profile in both the upper and lower channel as both have the same inlet volumetric flow rates.

**Figure 5:**
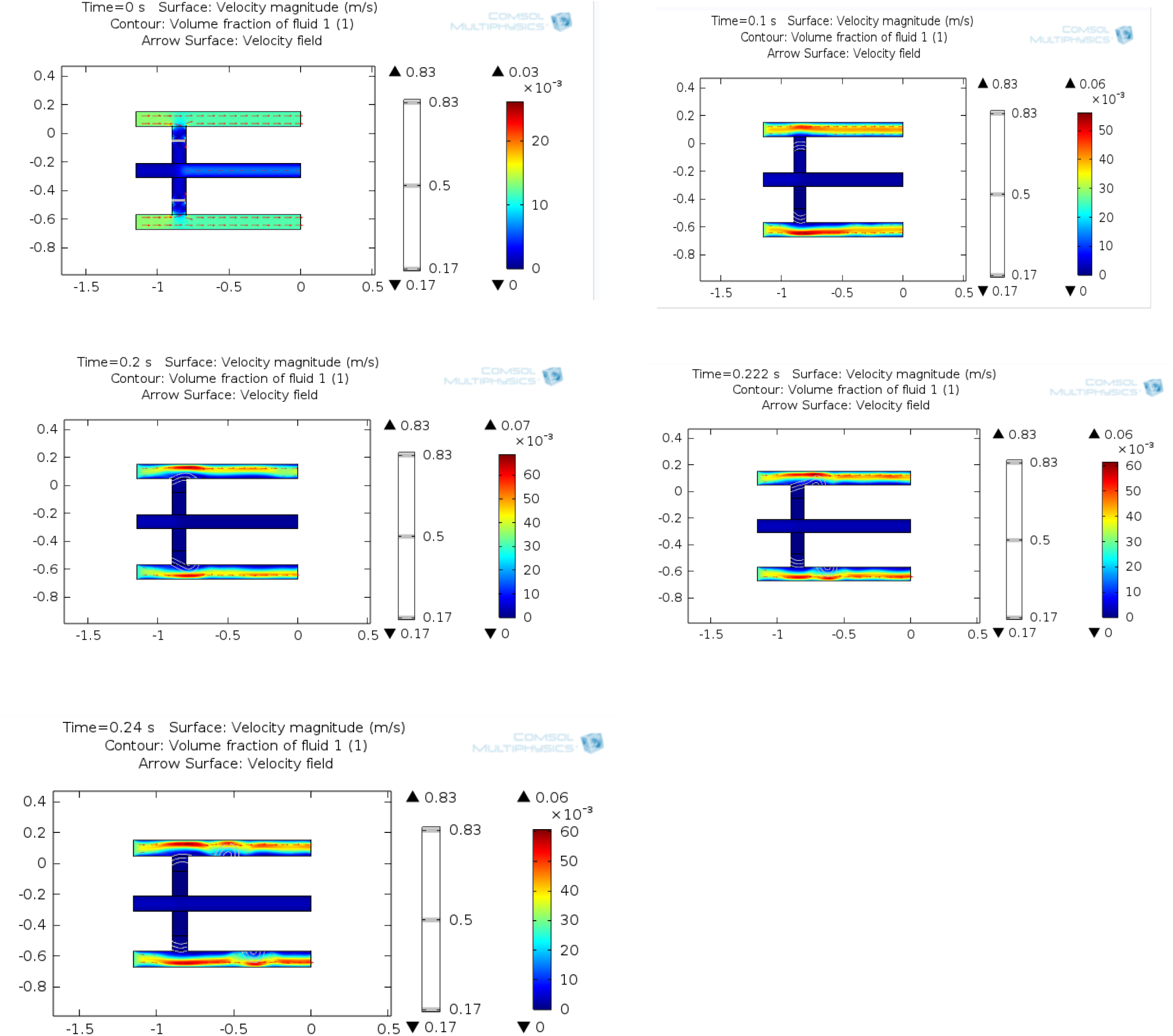
Velocity Profile of an E-Junction device.

## 4. Conclusions

The numerical simulation of the liquid-liquid two-phase droplet flow in an E-junction microchannel was investigated using Level Set Method (LSM) with COMSOL Multiphysics software. The droplet splitting was studied and the effective droplet diameter was reported.

Upon studying the E-Junction device it can be inferred that droplet splitting is taking place in both the channels in the laminar regime. As both the channels are of the same dimensions velocity throughout the process is the same which has been inferred from a velocity profile study. Droplet splitting is taking place following the process of filling up followed by necking then with pinch off and finally droplets are achieved. This process is continued till as long as there is a steady flow at the inlets.

The E-Junction device was developed to study droplet production in three phases as that is something another commonly used cross-flow device T-Junction was lacking. Further investigations are required to study such type of three-phase flows. The future scope of the work is in creating device configurations that will lead to the formation of distinct droplets in two or three-phase flow within a microfluidic chip that has a small footprint.

## References

Pingan Zhu; Liqiu Wang. Passive and active droplet generation with microfluidics: a review. Lab on a Chip. DOI:10.1039/C6LC01018K.

A. Abrishamkar; A.S. Rane; K.S. Elvira; R.C.R. Wootton; T. Sainio; A.J. DeMello. A COMSOL Multiphysics® Model of Droplet Formation at a Flow Focusing Device. COMSOL Conference at Rotterdam, The Netherlands.

Olsson, E, Kreiss, G, Zahedi, S. A conservative level set method for two phase flow II. J Comput Phys 2007; 225: 785–807.

Lei Lei; Hongbo Zhang; Donald J. Bergstrom; Bing Zhang; and Wenjun Zhang. Modeling Of Droplet Generation by a Modified T-Junction Device Using COMSOL. Applied Mechanics and Materials Vol 705 (2015) pp 112–116

Ich-Long Ngo; Trung-Dung Dang; Chan Byon and Sang Woo Joo. A numerical study on the dynamics of droplet formation in a microfluidic double T-junction. Biomicrofluidics. DOI: 10.1063/1.4916228

Jayaprakash Sivasamy; Teck-Neng Wong; Nam-Trung Nguyen; Linus Tzu-Hsiang Kao. An investigation on the mechanism of droplet formation in a microfluidic T-junction. Springer-Verlag. DOI 10.1007/s10404-011-0767-8

